# Microhomology-Mediated Tandem Duplication Drives Tandem Repeat Formation Across Life

**DOI:** 10.64898/2026.02.28.708716

**Authors:** Xianfang Wei, Wanxin Gong, Yifan Zheng, Jing Zhang, Xianyuan Wei, Chen Peng, Xiangwei He, Chao Jiang

## Abstract

Tandem repeats (TR) are common genomic elements that are highly variable and with major functional consequences. Yet, the evolutionary trajectory for their formation remains poorly understood. One proposed mechanism is microhomology-mediated tandem duplication (MTD), in which single-copy DNA segments flanked by microhomology undergo tandem duplication (TD) and can further expand into TRs. Although MTD was first identified in the fission yeast *Schizosaccharomyces pombe*, its universal occurrence and postulated role in TR evolution have not been established. Using whole-genome deep sequencing and new analytical tools, we show that MTDs occur *de novo* universally across bacteria, archaea, fungi, and viruses. Further analysis of 2,245 reference genomes and millions of isolate genomes from 103 prokaryotic and eukaryotic microbial species, combined with human population TD data, somatic-germline mutations, and disease-associated variants, reveals that MTDs are consistently the dominant TD-forming mechanism across domains of life. Evidence suggests that MTDs have initiated the formation of the majority of existing TRs in genomes. Importantly, MTDs also prevail in human pathogenic TR mutations, including those linked to cancers. Mechanistically, deletion of the conserved mutator gene Rad27 specifically increased *de novo* MTD frequency in the budding yeast *Saccharomyces cerevisiae*, implicating Rad27-mediated Okazaki fragment maturation in MTD formation. These findings establish MTD as a universal and functionally significant mechanism for TR genesis.

**Graphic abstract:** 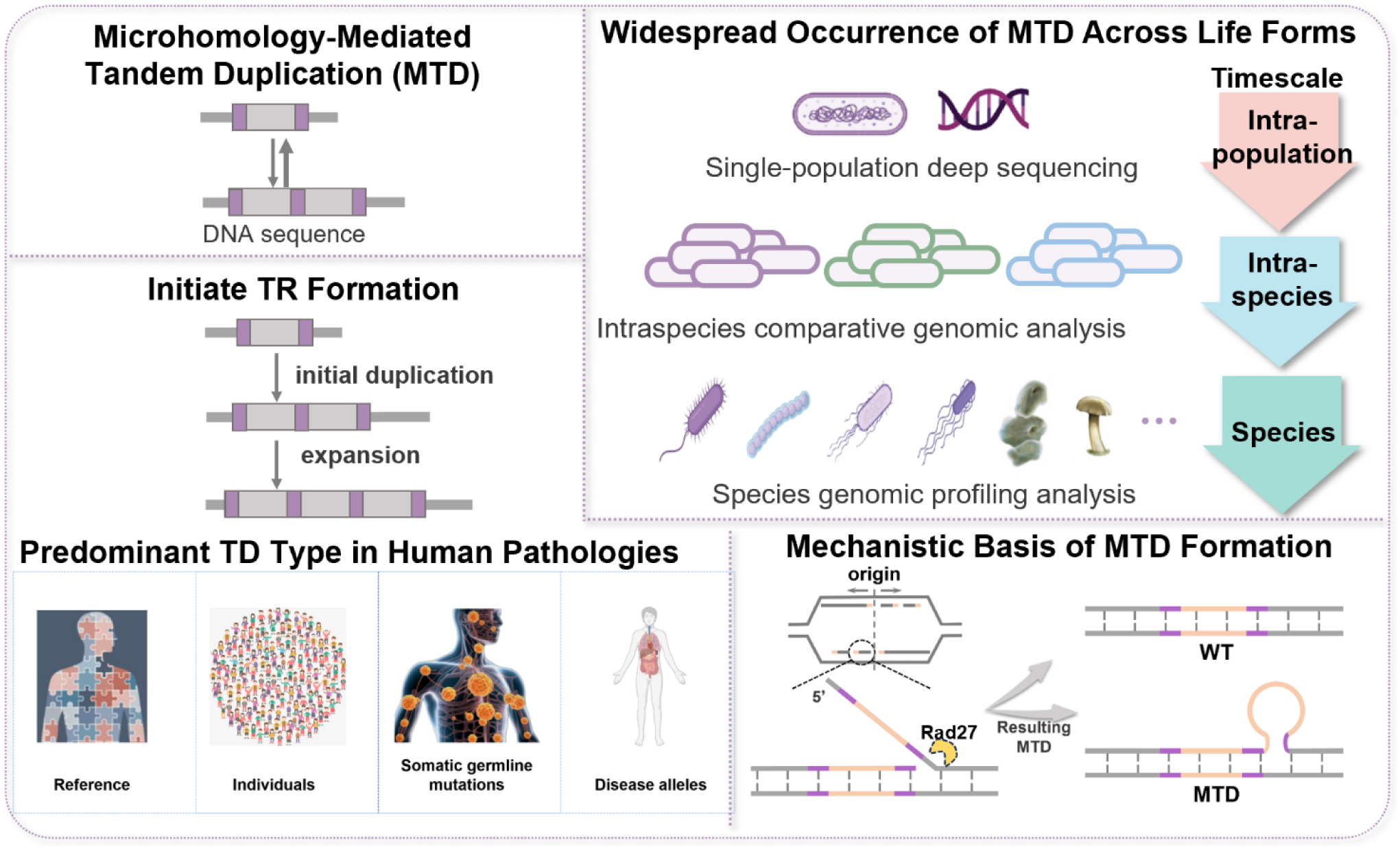

**Significance Statement:** Microhomology-mediated tandem duplications (MTDs) represent a universal mechanism driving tandem repeat formation across all domains of life—from viruses to humans. These duplications initiate genome evolution by expanding into functional tandem repeats and are the predominant form of pathogenic tandem duplications in human cancers and heritable disorders. Critically, disruption of the conserved Okazaki fragment processing pathway promotes MTD formation, establishing a fundamental link between DNA replication fidelity and genomic plasticity.

## Introduction

Repetitive sequences, regardless of the length of the repeat unit, exhibit high variability in copy numbers, making them key contributors to genomic instability. Repetitive sequences constitute a substantial portion of the genomes, more than 50% of the human genome, approximately 6% of which are in the form of tandem repeats (1) (2). Short tandem repeats with the length of the repeating unit typically ranging from two to 100 base pairs, are closely linked to various biological processes. For example, recent large-scale studies have revealed a genome-wide spectrum of tandem repeat expansions in humans, shedding light on their roles in human genetic diversity and diseases (2). Tandem repeats influence gene regulation, chromosomal architecture, and genome plasticity while also contributing to genomic instability, such as the formation of fragile chromosomal sites (3). Their instability is also implicated in the onset of complex diseases, including cancers and neurodegenerative disorders (4). Given their prominent role in shaping genome integrity and their association with diseases, tandem repeats are crucial for understanding the mechanisms of genetic variation, adaptation, and disease pathogenesis, as well as for gaining insights into genome evolution (2, 4).

The formation of tandem duplication (TD), signifying the transition from the unique sequence of a single-copy unit (1x) to the duplicated sequence of a doublet (2x), marks the inaugural step in the genesis of repetition in tandem repeats (TR). However, why tandem repeats appear at the specific loci in the extant genomes and how the specific DNA motives expand their copy number to various levels – in other words, the evolutionary origin for tandem repeats – remains elusive. Based on the sequence features of all tandem repeats in the genome of the budding yeast *Saccharomyces cerevisiae*, Haber and Louis (5) proposed that tandem repeats evolved from non-repetitive DNA segments with microhomology at the ends. Specifically, replication slippage between the microhomologous sequences flanking a short DNA segment or unequal crossing-over between them could initiate tandem duplication of the DNA segment, which may further expand into tandem repeats. However, despite over two decades of research, direct experimental evidence supporting this model remains limited.

In our previous work with the fission yeast *Saccharomyces pombe*, we identified a novel mutation form named Microhomology-mediated Tandem Duplication (MTD) (6). In MTD, single-copy short DNA segments with microhomology arms (MHAs) at the ends undergo tandem duplication, expanding from 1x to 2x. These duplicated DNA segments readily revert back into single copies, with the reversion frequencies more than ten times higher than those of *de novo* MTD formation measured at specific loci. Neither the total length (ranging from approximately ten to 500 base pairs) of the DNA segment nor the length of the microhomology arms (as short as four bps, by our working definition) impose stringent requirements for duplication. Millions of candidate MTD sites are found in the *S.pombe* genome. Indeed, MTD events arise broadly throughout the genome during cell proliferation. In a whole-genome deep sequencing analysis (10,000X coverage) of a single-cell-origin population of fission yeast cells, nearly 6,000 loci were identified where one or more sequencing reads carried an MTD, revealing unexpectedly high genetic diversity in a presumed isogenic population. However, at each locus, MTD-bearing cells are sparse, likely due to drift, negative selection, and the high reversibility of MTDs. This makes them difficult to detect in whole-genome sequence analysis using conventional analytical pipelines.

Importantly, *de novo* MTD exhibits high reversibility similar to the extant tandem repeats in the genome, but with a key difference that the latter are fixed at specific loci, whereas *de novo* MTDs can arise at numerous loci throughout the genome. It’s conceivable that the reversibility of MTD, combined with its prevalence throughout the genome, could provide a versatile mechanism for generating genomic plasticity, facilitating rapid adaptation in response to environmental changes (6). However, the prevalence of MTDs in other organisms and the molecular mechanism underlying MTD formation remain to be investigated.

The present study aims to elucidate the presence and traits of MTDs across all domains of life and explore their potential role in the evolutionary initiation of tandem repeats and their functional implications. We reason that MTD should be investigated at three different levels, corresponding to distinct stages along the hypothetical evolutionary trajectory of tandem repeats: (1) intra-population MTDs, referring to MTDs that arise *de novo* in a single-ancestor-originated cell population (*e.g.*, a single-colony population in microbes); (2) intra-species MTDs, referring to tandem duplications with MHAs found in the genomes of individual strains of the same microbial species, which likely represent fixed MTDs only in distinct strains/cell populations; and (3) pan-species MTDs, referring to tandem duplications with MHAs found in the reference genomes of various species, that are likely shared among all individuals of the species. Likewise, in metazoans such as humans, sporadic MTD variations among normal somatic cells within an individual can be considered *de novo* MTDs, while disease-associated MTDs — such as those found in tumors — would represent MTDs fixed in cancer cells due to pathological conditions.

Our pan-domain analysis across three evolutionary levels provides strong evidence that MTDs are widespread and are the dominant form of TDs, playing a pivotal role in genome evolution by initiating the formation of tandem repeats. They also serve as the raw materials for generating genomic plasticity that drives functional divergence and pathogenesis in prokaryotes and eukaryotes. Furthermore, our mechanistic studies in *S. cerevisiae* suggest that Okazaki fragment maturation during DNA replication, an evolutionarily highly conserved process, is closely associated with *de novo* MTD formation.

## Result

### Prevalence of *de novo* MTDs across domains of life

To assess the prevalence of *de novo* MTDs at the intra-population level, we conducted deep whole-genome sequencing (>1,000x coverage) in seventeen microbial organisms, including bacteria, archaea, and fungi, using the next-generation sequencing (NGS) platform (DNBSEQ-T7) on wild-type populations growing under no artificial selections (Table S1).

Using the established MTDfinder tool (details in Methods), we first identified a set of candidate MTD sites (*a.k.a.*, microhomology arms - MHAs, ranging from 4 to 25 base pairs)(6) in the reference genome for each species. The candidate MTD sites were found to be densely distributed and widely prevalent across different species. Notably, the number of these sites exhibited a linear correlation with the genome size (Figure 1A).

**Figure 1.**
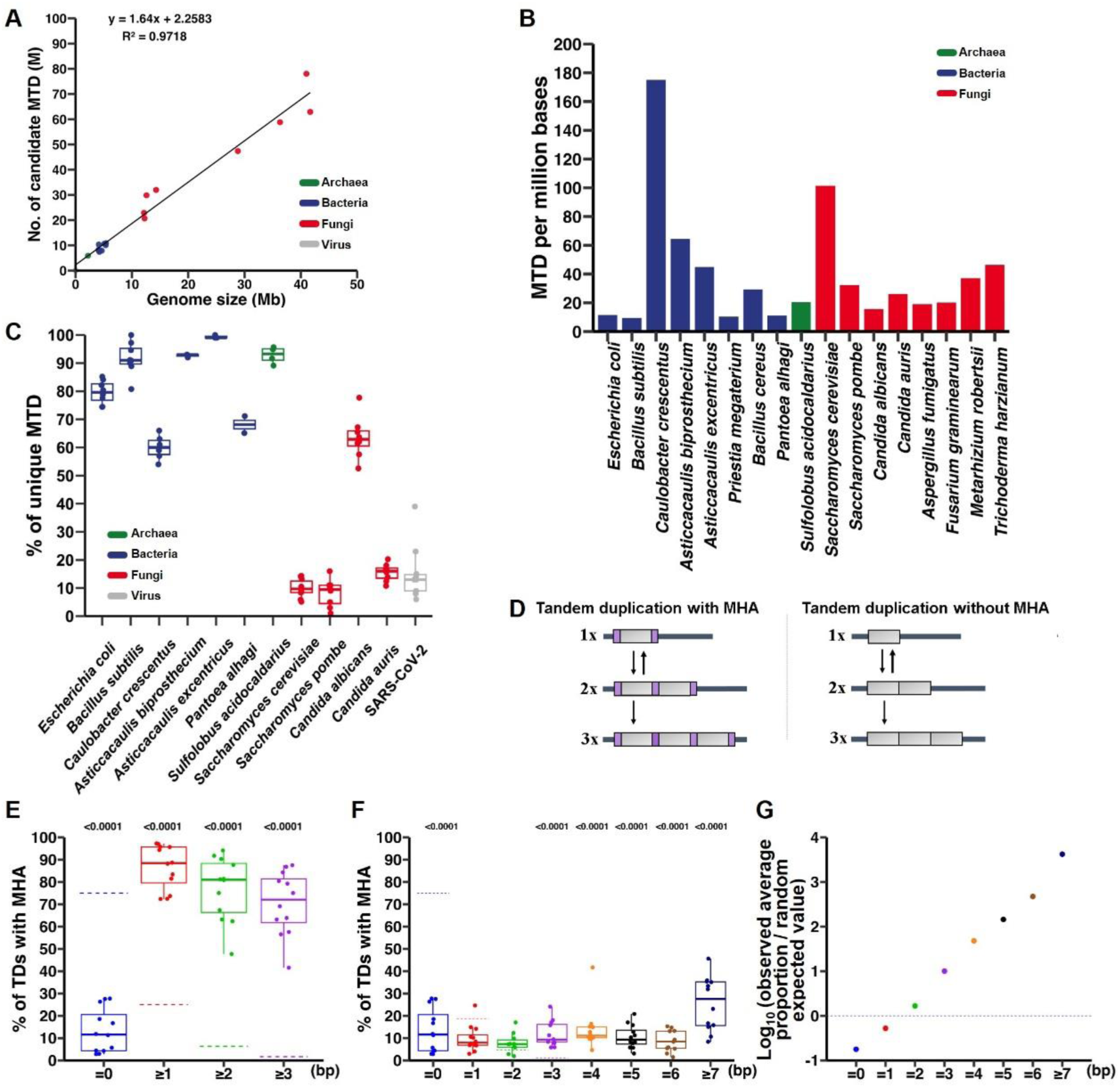
Genome-wide characterization of *de novo* MTDs across domains of life. **(A)** The relationship between genome size and candidate MTD site abundance across 18 species spanning fungi (red), bacteria (blue), archaea (green), and one virus (gray; species in Table S1). A unified regression line highlights a strong positive correlation (*R*² = 0.9718). **(B)** Frequency of *de novo* MTD loci per million bases across fungi (red), bacteria (blue), and archaea (green). MTD loci were identified using MTDfinder. **(C)** Proportion of unique MTD loci across population sequencing replicates for 12 species. Each point represents a replicate. Colors denote taxonomic groups: fungi (red), bacteria (blue), archaea (green), and viruses (gray). **(D)** A schematic representation of tandem duplication with and without MHA, with purple boxes indicating the microhomology arms. **(E)** Proportion of TDs with MHA in population-deep sequencing data. The x-axis represents MHA length (bp). Each point represents a species. Dashed line: random expected value. *P-values* were calculated using permutation tests (one-tailed, 100,000 simulations) against theoretical expectations. **(F)** Observed proportions of MHAs are categorized into eight length groups (0 bp, 1 bp, 2 bp, 3 bp, 4 bp, 5 bp, 6 bp, and ≥ 7 bp) and compared to random expectations (dashed line). Each point represents a species. *P*-values were calculated as in (E). **(G)** The log_10_-transformed ratios of the observed average proportions of TDs at different MHA lengths (from Figure 1F) compared to their corresponding random expected values. Each point represents the logarithmic transformation of the ratio for a specific MHA length, highlighting the discrepancies between actual observations and random expectations.

For each MHA pair, a unique probe sequence was created to match the junction region of the putative tandem duplication formed by the corresponding DNA segment. *De novo* MTD mutations were then detected in deep sequencing datasets by identifying DNA reads carrying an insertion that matched with the probe sequence. The results revealed the universal presence of *de novo* MTDs in all the examined species, with varying frequencies of MTDs, ranging from 6.77 to 104.67 per million bases per thousand-fold genome coverage (Figure 1B). Notably, by analyzing the published sequencing datasets from ten viral species with a DNA or RNA genome, we detected high frequencies of MTDs ranging from 17.51 to 261.73 per million bases per thousand genome coverage (Table S2). We suggest that *de novo* MTD is also conserved in DNA or RNA viruses and that high depth of whole-genome sequencing (commonly associated with viral genome entries in databases), overall, is conducive to MTD detection.

We further verified *de novo* MTD mutations using alternative methods. First, we selected a series of high-frequency MTDs in *Escherichia coli (E. coli)* and *S. pombe*, through PCR amplification followed by Sanger sequencing, successfully identified the DNA segments harboring MTDs (Figure S1), supporting the reliability of the current sequencing and analysis pipeline for MTD detection. In addition, we employed an alternative sequencing platform (Illumina NovaSeq 6000) for *S. cerevisiae*, which demonstrated minimal differences in MTD detection compared to the primary sequencing platform (DNBSEQ-T7). Furthermore, single-cell whole-genome DNA sequencing in *S. cerevisiae* revealed that the proportion of MTDs was highly consistent with population-level NGS data (Table S3; Methods). These results demonstrate that the MTDs identified across various species using the established procedure are genuine overall.

Next, we compared the detected MTD sites across independent biological replicates, revealing species-specific features in MTD consistency (Figure 1C). In some yeast species, such as *S. cerevisiae* and *S. pombe*, 90% of MTD sites were consistent across biological replicates, indicating that the locations and frequencies of the MTDs are highly consistent in these species. *Candida albicans* exhibited a unique MTD percentage of approximately 60%, and *Candida auris* had a unique MTD percentage of a low 10%. However, in four bacterial species, *E. coli*, *Bacillus subtilis, Asticcacaulis biprosthecium,* and *Asticcacaulis excentricus*, each replicate had mostly unique MTD sites, with only 10% of MTD sites shared between replicates. These findings suggest that the patterns of MTD occurrence may vary significantly among different species.

In summary, population deep-sequencing analyses across various microbial species, including prokaryotes, eukaryotes, viruses, and archaea, confirmed the prevalence of *de novo* MTDs in all tested species.

### MTD is the dominant form of *de novo* tandem duplication events

Tandem duplication (TD) may occur with or without MHA (MTD or NMTD, respectively) (Figure 1D). Because MTDfinder depends on the presence of MHA for MTD calling, NMTD identification is precluded. To assess the importance of MHAs in TD formation, we developed MTDfinder-pro to simultaneously identify MTDs and NMTDs by directly detecting TDs in individual reads, followed by assessing the presence of flanking MHAs (Method). It is important to note that this method is constrained by the read lengths used for analysis (150 bp in this study), which likely results in an underestimation of the actual number of TDs. Nonetheless, cross-species analysis of whole-genome sequencing datasets for fifteen species using both MTDfinder and MTDfinder-pro revealed perfect concordance in MTD loci identification. All 652 overlapping candidate loci were consistently detected by both algorithms with 100% agreement, demonstrating excellent reliability and robustness of the MTD detection pipeline.

Using MTDfinder-pro, we analyzed deep sequencing datasets and calculated the MTD/TD ratio (Figure 1E). Given the roughly equal presence of four nucleotide bases, the probability of at least one bp being identical by random on both ends of a duplicated segment is 25%. In other words, loci without MHAs (0 bp) should account for 75% of all tandem duplications, whereas the expected random occurrence of loci containing MHAs longer than three bp is about 1.6% (1/64). However, among TDs detected in twelve bacteria and fungi, the proportion of loci with MHA ≥ 3 bps was significantly higher, constituting 70.41% of the total TDs (p < 0.0001, permutation tests against theoretical expectations). Conversely, the proportion of loci with MHA = 0 bp was significantly lower than expected, accounting for only 13.32%. Consistent results were obtained across multiple sequencing replicates for each species (Figure S2). These findings demonstrate that TDs without MHAs do occur *de novo,* although TDs with MHAs - especially those of ≥ 3 bps - significantly exceed random expectations, representing the dominant form of TDs in the examined species.

We further examined the distribution of MHA lengths in tandem duplication events across ten species with sufficient MTDs detected and compared these results to the expected random values (Figure 1F). We observed a clear trend - longer MHA lengths correlated with greater positive deviations from random background expectations (Figure 1G), indicating that longer MHAs are preferably involved in generating tandem duplications.

In summary, at the intra-population level, the presence of MHAs in *de novo* TDs is dominant, with longer MHAs more favored, underscoring their significance in generating TDs.

### MTD is the dominant form of TDs at the intra-species level

The MTDs characterized above were generated *de novo* within an isogenic cell population. We reckoned that these *de novo* MTDs provide the raw materials upon which evolutionary forces can act. Under proper conditions, specific MTDs would be fixed within the population. To test this idea, we sought to determine whether MTDs remain the dominant form of TDs at the intra-species level.

Newly fixed MTDs in a species should be found in some cell populations but not others. Because the genome assemblies found in public databases typically report the major haplotype of a strain, we reasoned that the TDs uniquely identified in a specific genome assembly could be considered “fixed” within the corresponding cell population. This contrasts with *de novo* MTDs, which are typically present only in very small sub-populations.

We sought to identify such TDs by analyzing 1.5 million independent genomes extracted from NCBI databases, covering 103 species, including sixty-six bacterial, thirty-two fungal, and five archaeal species, each with a sufficiently high number of genome entries (Methods). This analysis involved aligning all the genomes for each species to identify genome-specific TD loci and excluding those that were too short or exhibited low complexity in sequence (Methods). In total, ∼76,000 loci were identified, referred to as “variable TD” loci, as they were found in some but not all genome entries within a species. These loci represent TDs fixed specifically (and thus recently) in distinct cell populations. The distribution of the number of genomes and variable TD loci among the 103 species is shown in Figure 2A. In bacteria, *E. coli* exhibits the highest number of variable TD loci, reaching approximately 13,000, while in fungi, *S. cerevisiae* has the highest number, reaching approximately 9,000 (Figure 2A). Notably, more genomes do not always lead to more variable TD loci.

**Figure 2.**
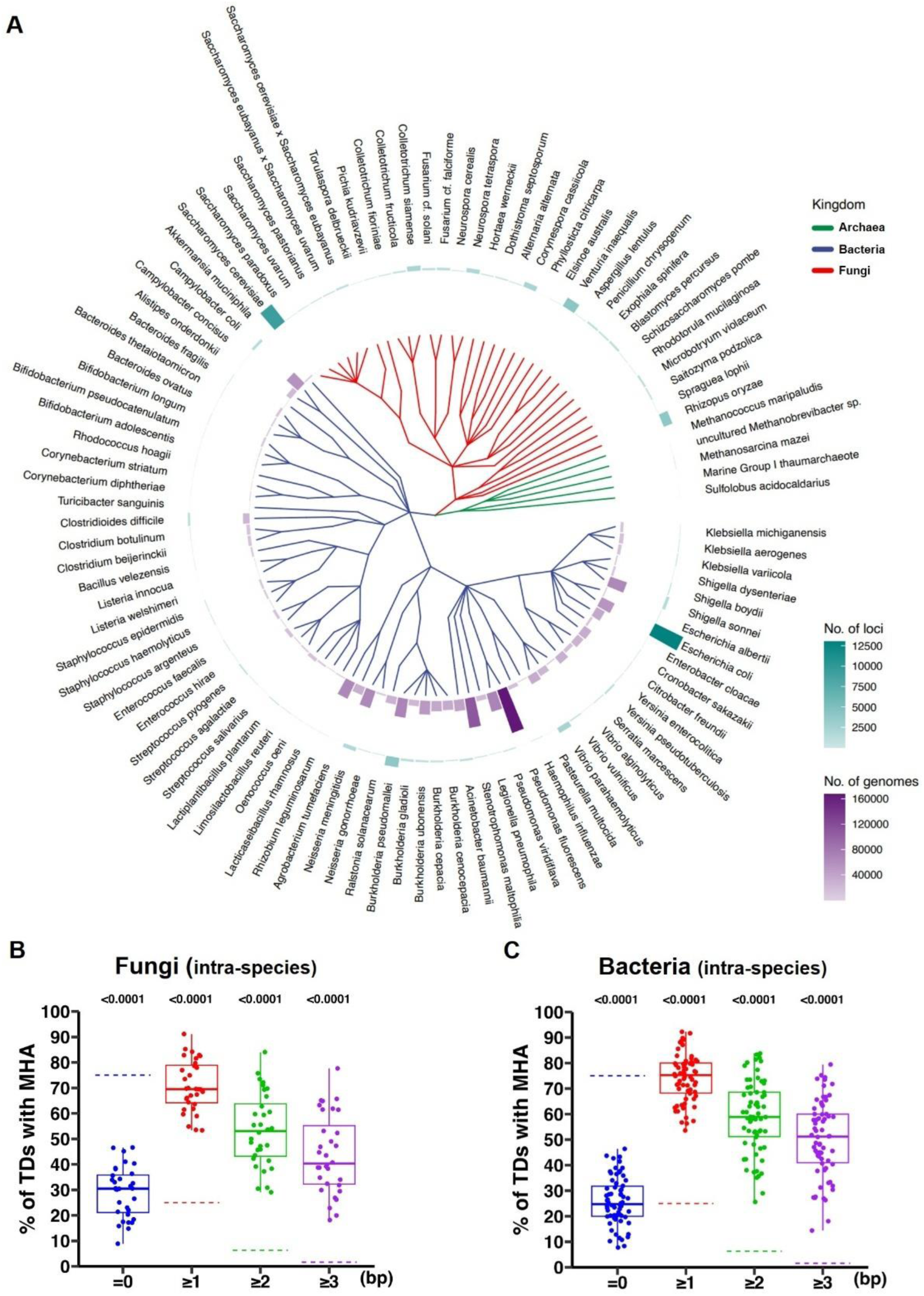
MTD is the dominant form of variable TDs at the intra-species level. **(A)** Distribution of the number of genomes and variable TD loci across the 103 species. The green bars represent the number of genomes, while the purple bars indicate the number of variable TD loci. **(B-C)** Proportions of variable TDs with MHA at the Intra-species level of fungi (B) and bacteria (C). Each point represents a species, and the dashed line represents the random expected value. *P-values* were calculated using permutation tests (one-tailed, 100,000 simulations) against theoretical expectations. Data are presented for species with at least 20 TD loci.

Among these variable TD loci across bacteria, fungi, and archaea, consistent patterns emerged: only approximately 25-30% of the variable TD loci lacked microhomology arms, significantly lower than the expected random value of 75%. Moreover, about 50% of the variable TD loci contained MHAs longer than 3 bps, far exceeding the expected random proportion of 1.6% (Figure 2B, 2C, Figure S3). These results demonstrate that fixation of tandem duplication is actively and prevalently ongoing in extant microbial populations. Furthermore, MTD is also the preferred form of TD at the intra-species level, consistent with the notion that *de novo* MTDs may persist and prevail among the populations once generated.

### MTD is the dominant form of TDs at the species level

We further hypothesized that once fixed in distinct cell populations or natural isolates, some MTDs could persist at long evolutionary time scales, which may be detected in the reference genomes of all species.

To test this, we quantified the proportion of MTDs among all TDs across approximately 4,000 species spanning different domains of life. To ensure diversity, we selected 1,000 representative genomes per kingdom, excluding species with fewer than ten TD loci. Our analysis, which included fungal, protozoan, bacterial, and archaeal species, revealed that approximately 50%-60% of TDs lacked MHAs (Figure 3A-D), significantly lower than the random expectation value (75%) but significantly higher than those at the species level (Figure 3E-H). We reckoned that this might be due to the homology decay of the nascent MHAs over a longer evolutionary time scale at the species level. To evaluate this possibility, we re-analyzed TDs with imperfect repeat unit matches, identified by the Tandem Repeats Finder program (7) with a match score greater than 30 (see details in Methods). By including these TDs that may have experienced sequence decay, the percentage of TDs without MHAs was drastically reduced to around 20% across all species analyzed (Figure 3I), indicating the MTD proportion returns to 80%—a value similar to the proportions observed at the intra-species level (Figure 3J). This finding suggests that, as time progresses, the original MHA may become degenerate, akin to the degeneration of the repeat units seen in the inactive CRISPR array systems (8).

**Figure 3.**
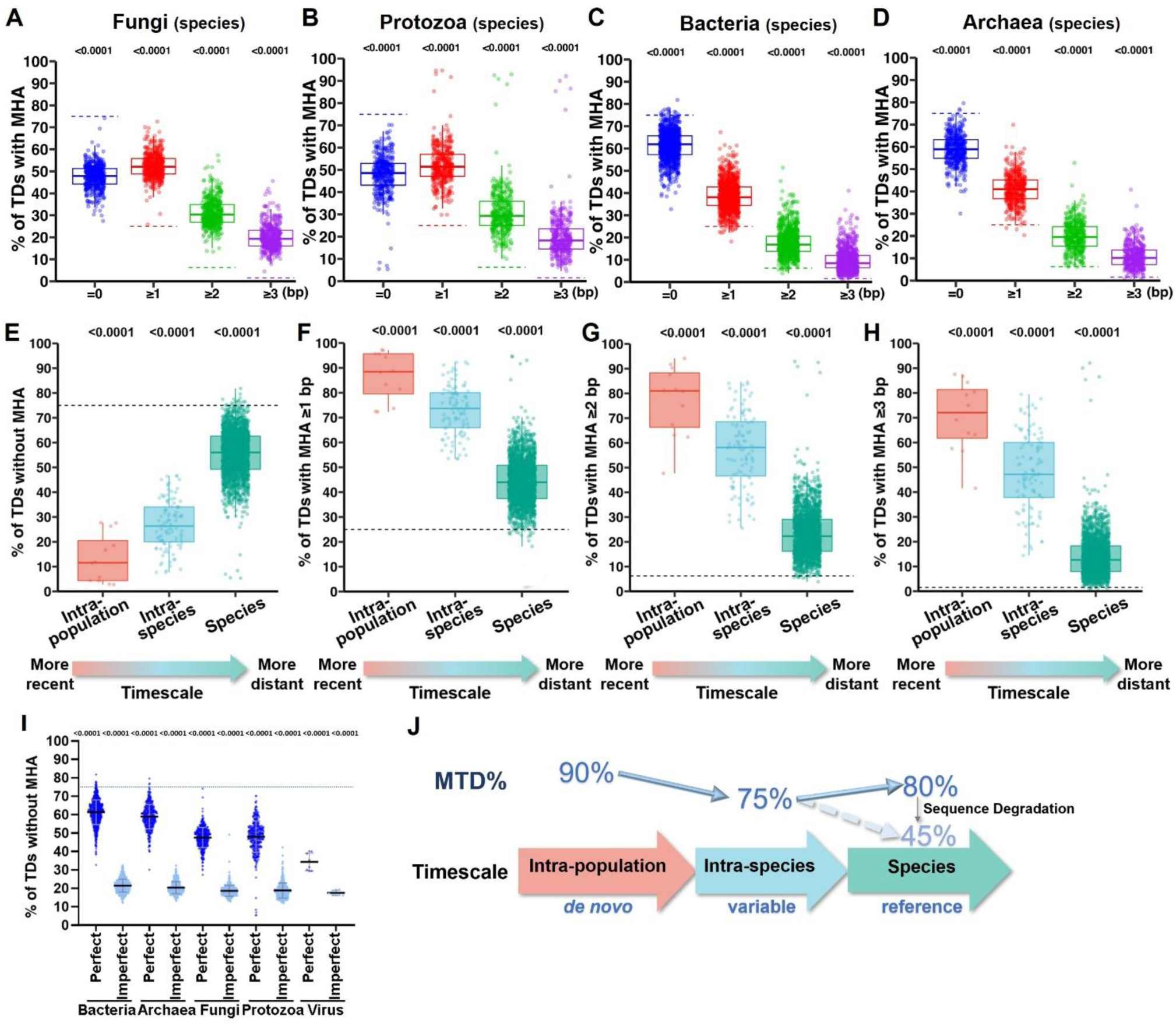
MTD is the dominant form of TD across evolutionary time scales. **(A-D)** Proportions of TDs with MHA at the species level of fungi (A), protozoa (B), bacteria (C), and archaea (D). Each point represents a species, and the dashed line represents the random expected value. *P-values* were calculated using permutation tests (one-tailed, 100,000 simulations) against theoretical expectations. Data are presented for species with at least 10 TD loci. **(E-H)** Proportions of TDs without MHA (E), MHA ≥ 1bp (F), MHA ≥ 2bp (G), and MHA ≥ 3bp (H) across evolutionary scales. Intra-population level (red), intra-species level (blue), and species level (green). The timescale is shown at the bottom, with the direction of the arrow indicating increasing evolutionary distance. The dashed line represents the random expected value. *P-values* were calculated using permutation tests (one-tailed, 100,000 simulations) against theoretical expectations. **(I)** Proportions of TDs without MHA across various phylogenetic kingdoms, categorized by the degree of mutation accumulation. Dark blue bars denote the ‘Perfect’ group, where unit match percentage equals 100%, and light blue bars represent the ‘Imperfect’ group, with unit match percentage below 100%. **(J)** Schematic illustrating the relationship between evolutionary time and MTD proportions: Intra-population (short-term): 90% MTD frequency (*de novo* mutations); Intra-species (mid-term): 75% MTD (polymorphic loci in isolates); Species (long-term): MTD% increases to 80% after taking sequence decay into consideration (match score >30).

### MTDs may further expand to form multi-copy tandem repeats

Haber and Louis proposed that the formation of TD (1x -> 2x) serves as the initiation step for the formation of multi-copy tandem repeats (TRs) (Figure 4A). Having established the occurrence of *de novo* duplications, we next asked whether the MTDs within the population characterized above may undergo further expansion to form TRs as proposed. If so, the variable TD loci identified earlier (Figure 2B-C) might include certain loci that show high variability in copy number. To test this hypothesis, we examined 103 species where reference genomes showed single-copy (1x) loci, and natural isolates exhibited at least three TD copy number variants (predominantly 1x with minor 2x/3x cell populations, Figure 4B). These highly variable TD loci may represent active sites of TR expansion.

**Figure 4.**
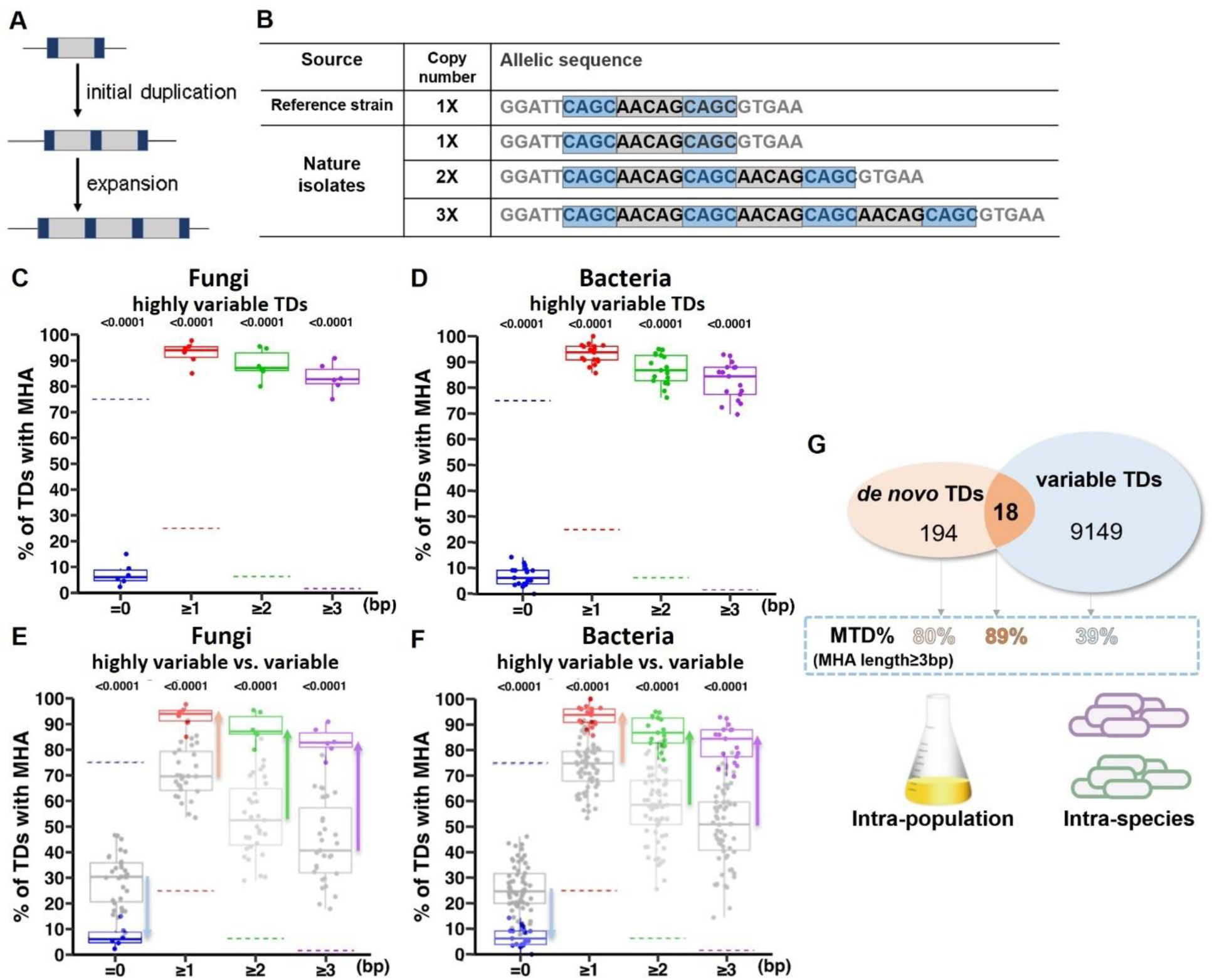
MHA drive the expansion of TD into TR. **(A)** A model illustrating the origin of tandem repeats. Gray boxes represent the core region of a tandem repeat, while blue boxes indicate microhomology arms. **(B)** Schematic of highly variable TDs, using an example locus in *E. coli* (NC_000913.3:4593203-4593225). **(C-D)** Proportions of TDs with MHA at the highly variable TDs of fungi (C) and bacteria (D). Each point represents a species, and the dashed line represents the random expected value. *P-values* were calculated using permutation tests (one-tailed, 100,000 simulations) against theoretical expectations. Data are presented for species with at least 20 TD loci. **(E-F)** Comparative analysis of MHA-containing TDs in highly variable versus variable TDs across fungi (E) and bacteria (F). Gray plots: variable TDs; colored plots: highly variable TDs. Arrows represent the percentage difference, categorized by: no MHA (blue), MHA ≥1 bp (red), MHA ≥2 bp (green), and MHA ≥3 bp (purple). Statistical parameters as in C-D. **(G)** Overlap analysis of *de novo* TDs at the intra-population level and variable TDs at the intra-species level identifies putative fixation intermediates. Blue inset: proportion of TDs with ≥3-bp MHAs in *de novo*, overlapping and variable TDs.

Among all 103 species examined, highly variable TD loci were identified, ranging from a few dozens to a few hundreds, with *Burkholderia pseudomallei* exhibiting the highest number (N=485) (Table S4). The number of highly variable TD loci in archaea was too low to support statistical analysis, possibly due to the limited number of genomes per species. Interestingly, in both bacteria and fungi, over 90% of highly variable TD loci harbored MHAs (Figures 4C-D)—a proportion significantly exceeding the MTD frequency in conventional variable TD loci (Figures 4E-F). Together, these findings support the notion that the evolution of tandem repeats—from a single copy (1x) to multi-copies (2x, 3x or higher)—is preferentially mediated by MHA.

Furthermore, in *S. cerevisiae*, we analyzed overlapping TD loci from two sources: *de novo* TDs identified in a laboratory standard strain cell culture at the intra-population level and variable TDs identified among natural isolates at the intra-species level. These TD loci may be always active to generate genomic diversity. Notably, all overlapping TD loci were MTDs, with 90% containing MHAs ≥3 bp, significantly exceeding random expectations (*P* < 0.0001), the *de novo* MTD proportion (80%), and the MTD proportion in variable TD loci (39.2%) (Figure 4G). More important, among these overlapping loci, 88.3% (5/18) were highly variable TDs, versus only 1.4% (131/9,149) in the broader variable TD loci pool at the intra-species level (Table S5), indicating active TDs are likely mediated by MHA.

This multi-scale convergence at the intra-population, intra-species, and species levels demonstrates that MTDs formed in single-cell experiments evolve into natural polymorphisms, where they serve as evolutionary springboards for selection by various evolutionary forces and subsequent expansion into TR (Figure 4A, 4G). Thereby, we establish that MHAs not only facilitate TD formation but also govern their expansion potential.

### Abundant MTDs in the human genomes are associated with diseases

To investigate the prevalence and potential functional significance of MTDs in humans, particularly their association with diseases, we conducted a three-tiered analysis following the approach used in microbial studies. First, we identified tandem duplications (TDs, 2X; 423,331) and tandem repeats (TRs, 3X or higher; 4,719) in the human reference genome (hg38) using stringent criteria: TRs were defined as repeats with unit size ≥8 bp and 100% sequence identity. This conservative approach explains the lower TR count relative to conventional databases. Our analysis revealed that over 50% of the TDs and nearly 80% of the TRs in the human reference genome carry microhomology arms (Figure 5A), indicating that MTDs are conserved in the human genome and may represent a predominant class of tandem duplication. The increase MHA% in TR again indicates that MHA is more prevalent in more active TD loci.

**Figure 5.**
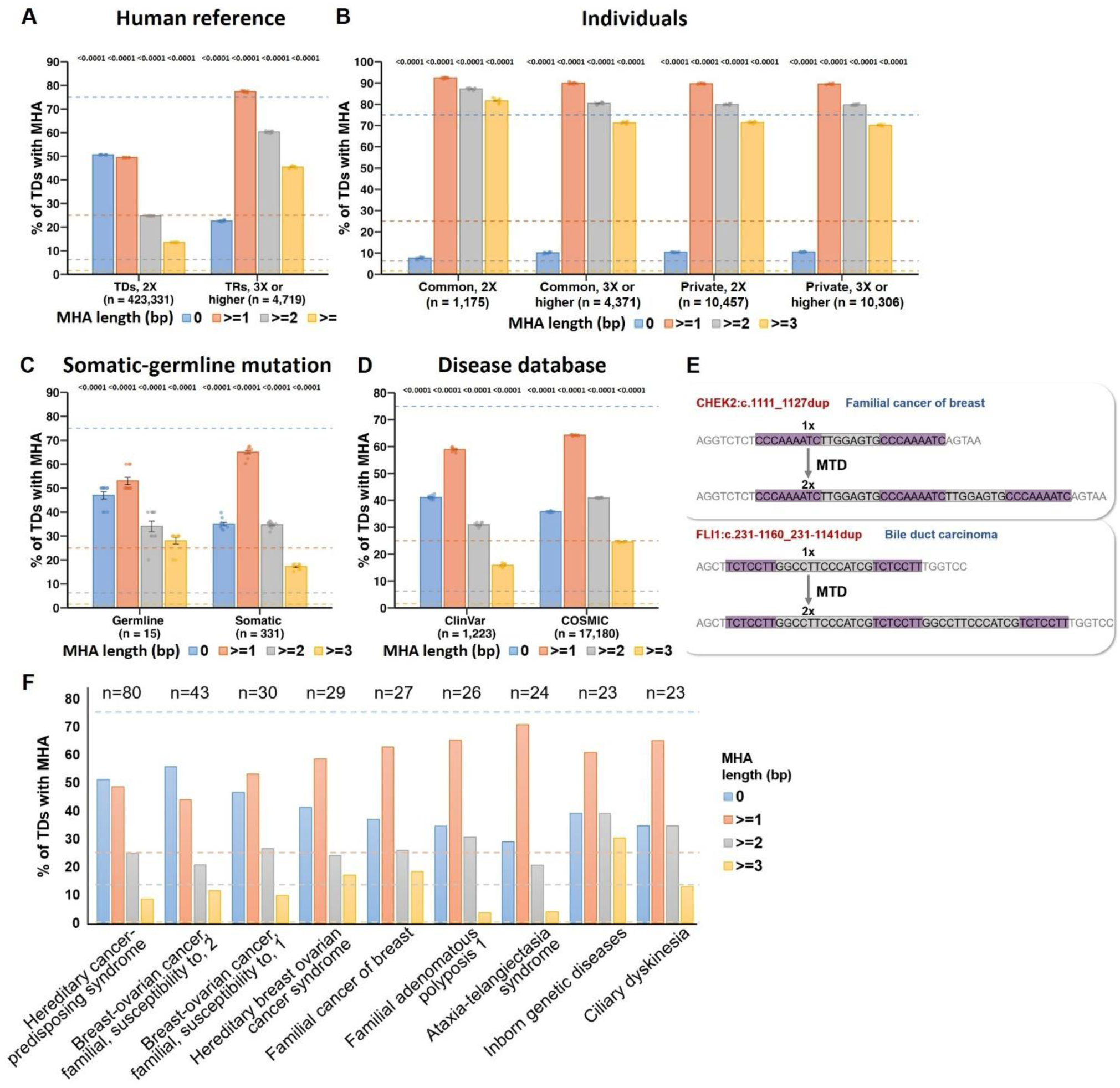
MTD analysis in the human genome and its associations with diseases. **(A)** Proportions of tandem repeats with MHA in the human reference genome. The dashed line represents the expected value based on random distribution. The left panel shows tandem repeats with a copy number of 2x (n=423,331), while the right panel shows tandem repeats with higher copy numbers (n=4,719). *P-values* were calculated using permutation tests (one-tailed, 100,000 simulations) against theoretical expectations. **(B)** Proportions of variable tandem repeats with MHA in 2,770 individuals. Common loci (n = 5,546): non-reference alleles occurred in at least 5% of a population, and private loci (n = 20,763) occurred in less than 1%. The dashed line represents the expected random value. The left panel shows variable tandem repeats (2X), while the right panel shows variable tandem repeats (3X or higher). *P*-values were calculated as in (A). **(C)** Proportions of TDs with MHA in somatic and germline mutations. The dashed line indicates the expected random value. The left panel represents somatic mutations (n=331), while the right panel shows germline mutations (n=15). *P*-values were calculated as in (A). **(D)** Proportions of TDs with MHA in mutations associated with human diseases. The dashed line indicates the expected random value. The left panel represents mutations from the ClinVar database (n=1,223), while the right panel shows mutations from the COSMIC database (n=17,180). *P*-values were calculated as in (A). **(E)** Example of MTD sequences from disease databases. The top sequence represents the reference (1x), and the bottom represents the mutated sequence (2x), with purple shading indicating the MHA. The first mutation is from ClinVar, located at CHEK2 (checkpoint kinase 2): c.1111_1127dup, associated with familial breast cancer; the second is from COSMIC, located at FLI1 (Fli-1 proto-oncogene): c.231-1160_231-1141dup, associated with Bile duct carcinoma. **(F)** Proportions of TDs with MHA in mutations across nine human diseases. The dashed line indicates the expected random value. Different colors represent different lengths of MHA, and “n” denotes the number of loci analyzed.

Next, we conducted inter-individual comparative genomic analyses on data from approximately 3,000 individuals (9), analogous to the intra-species analysis in microbes. This analysis revealed extensive variable tandem repeats, including 11,632 tandem duplications (2X) and 14,677 higher-order tandem repeats (3X or more). Among them, common loci (n = 5,546) were defined as non-reference alleles present in at least 5% of the population, while private loci (n = 20,763) were found in less than 1%. Notably, nearly 90% of variable tandem duplications and repeats carry MHAs, far exceeding the random expectation of 25% (Figure 5B). This starkly contrasts 50% MHA rate in the 2X TDs of the human reference genome (Figure 5A). The higher prevalence of MHAs at the inter-individual level supports a key role for MTDs at active TD loci, and older MHAs may be subjected to sequence decay.

Finally, we analyzed deeply sequenced somatic and germline genomes across multiple tissues from the same individuals(10), analogous to intrapopulation analysis in microbes. Among *de novo* TDs detected in laser-microdissected human tissues, 65.0% (215/331) of somatic and 53.3% (8/15) of germline events exhibited MHA (Figure 5C), providing strong evidence that MTD contributes to ongoing variation in the human genome.

Based on these findings, we hypothesized that MTDs may contribute to human diseases. To test this, we analyzed approximately 20,000 TD loci from the Cosmic Cancer and the ClinVar disease databases. Remarkably, 60-70% of these TD mutations are associated with MHAs (Figure 5D). Two examples—CHEK2 (checkpoint kinase 2): c.1111_1127dup, associated with familial breast cancer, and FLI1 (Fli-1 proto-oncogene): c.231-1160_231-1141dup, associated with bile duct carcinoma—illustrate disease-linked MTDs (Figure 5E), supporting a potential significant role of MTDs in human diseases.

Focusing on specific diseases, we analyzed TDs in nine disease types with more than 20 TD-containing alleles, including six cancer-related diseases and three genetic disorders (Figure 5F). In all cases, the proportion of TDs with MHA was significantly higher than expected by chance.

In summary, these results demonstrate that MTDs represent the predominant form of TDs in humans and are broadly associated with disease.

### Rad27-mediated Okazaki fragment maturation is a major MTD formation mechanism

Mutators are genes involved in DNA metabolism whose dysfunction drastically increases specific mutation frequencies genome-wide. Among them, Rad27 deficiency uniquely elevates insertion and deletion (indel) rates by over 100-fold in *S. cerevisiae* (11). Rad27, also known as FEN1 (Flap Endonuclease 1), is a Rad2 nuclease family member (12), essential for processing Okazaki fragments during DNA replication (13). We speculated that Rad27 may be implicated in *de novo* MTD formation.

Indeed, by re-examining the spectrum of isolated mutations in *rad27Δ* mutants in *S. cerevisiae* (14), we found that most insertion alleles contained MHAs, qualifying them as fixed MTDs within the population (Figure S4). We also re-analyzed the published whole-genome sequencing dataset for *S. cerevisiae* wild-type and *rad27Δ* mutants(15), using MTDfinder or MTDfinder pro to identify *de novo* MTDs. Despite the fact that the dataset was at relatively low coverage (250x), we found a significant increase in MTD density in *rad27Δ* (232 MTDs per million bases) compared to wild-type (122 MTDs per million bases) and *rad27Δ* revertants (98 MTDs per million bases) (Figure 6A). These results suggested that compromised Okazaki fragment processing may contribute to MTD formation, whether due to genetic perturbation (*rad27Δ*), external stresses, or replication errors.

**Figure 6.**
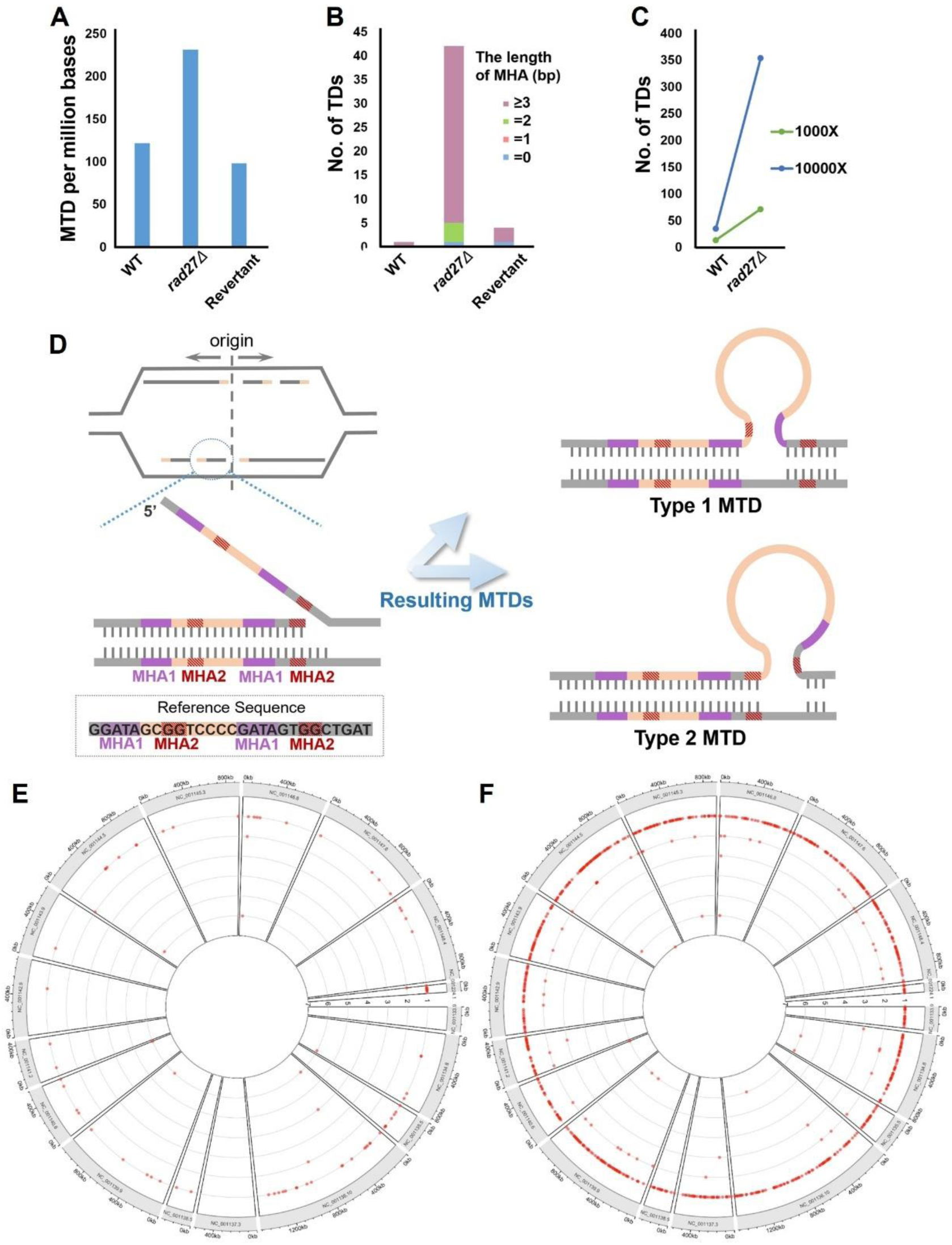
Rad27 deficiency promotes MTD formation in *S. cerevisiae*. **(A-B)** *De novo* MTD frequency and TD distribution. (A) Frequencies of *de novo* MTDs and (B) TD distribution with associated MHA lengths in WT, *rad27Δ*, and *rad27Δ* revertant strains. *S. cerevisiae* cells were grown at 30°C. MTD loci were identified through whole-genome sequencing (WGS) using MTDfinder (A) and MTDfinder-pro (B). **(C)** *De novo* TD counts at different sequencing depths. TD counts in WT and *rad27Δ* strains at 1,000× and 10,000× sequencing depths. The WT cells were cultivated at 30°C, while the *rad27Δ* strain was grown at 25°C. The TD loci were identified via WGS conducted on the DNBSEQ-T7 platform, utilizing MTDfinder-pro for the analysis. **(D)** Model of MTD generation during Okazaki fragment processing. Left panel: Schematic of replication origin dynamics (top) and MHA configurations. Type 1 MHA (purple) and Type 2 MHA (red-striped) are defined by distinct sequence alignments and flanking regions. Right panel: Resulting tandem duplication events. Type 1 MTD (top) arises from the pairing of Type 1 MHA, while Type 2 MTD (bottom) results from the pairing of Type 2 MHA, leading to the insertion of different sequences. **(E-F)** Genome-wide TD recurrence in **(E)** WT and **(F)** *rad27Δ*. Rings represent chromosomes and mitochondrial DNA; the y-axis shows TD recurrence in 10-bp windows.

To accurately assess the impact of *rad27Δ* on *de novo* MTD formation, we performed ultra-deep sequencing using the DNBSEQ-T7 platform, with coverage ranging from 1,000× to 10,000×, representing a 40-fold increase over the published dataset (15). Consistently, higher sequencing depth revealed up to 35 times more *de novo* TDs in the corresponding yeast strains. Notably, *rad27Δ* exhibited a 7-10-fold increase in *de novo* TDs compared to WT (Figure 6B-C). Notably, *rad27Δ* strains showed no significant difference in SNP burden compared to WT (mean 4,887 vs 5,367 SNPs; unpaired t-test, *P* = 0.79), demonstrating that Rad27 specifically modulates MTD formation.

Rad27 cleaves the 5′ RNA-DNA flap during maturation; in its absence, single-stranded flaps persist before other nucleases process them at low efficiencies (16, 17). We propose that the accumulated single-strand flaps may drive MTD formation, probably via a single-stranded DNA invasion process that involves microhomology-mediated annealing (Figure 6D). Considering the Okazaki fragments’ non-random distribution in the genome and their consistent average length (18), we hypothesized that if *de novo* MTDs are derived from the 5’ flaps of Okazaki fragments, their distribution should follow distinct patterns.

To this end, we compared the genome distribution of TDs among independent biological replicates of WT and *rad27Δ*, respectively. In contrast to the dense distribution of the 5’ flaps (average inter-flap distance ∼ 190 nt, estimated by electron microscopy)(18), *de novo* TDs were detected sparsely, even at 10,000× sequencing depth, with 124 in WT and 993 in *rad27Δ* (Figure 6E-F). Nonetheless, significant clustering of *de novo* TDs was detected, with 41.9% (52/124) of TDs in WT and 16.7% (166/993) in *rad27Δ* colocalizing within the same 10 bp window, far exceeding random expectations (P<0.001, binomial test). This may be partially explained by high consistency in replication origin utility and Okazaki fragment formation across different rounds of genome replication.

Strikingly, multiple clusters (19 in *rad27Δ* and 5 in WT) harbored overlapping MTDs, with sequencing alignment consistent with the scenario in which the same single-stranded DNA fragment undergoes alternative annealing using different microhomology arms (Figure 6D, S5). Together, these findings support the model that the persistent 5’ flaps promote microhomology-driven annealing, leading to MTD formation during error-prone ligation (Figure 6D).

## Discussion

Our analysis highlights the widespread occurrence of Microhomology-mediated Tandem Duplications (MTDs) across diverse organisms and their central role in genomic evolution, environmental adaptation, and human diseases.

### Evolutionary significance of MTD

MTDs uniquely generate widespread, reversible genetic variation *de novo*. Although most MTDs are neutral or disadvantageous and thus quickly reversed or eliminated, some duplications become fixed within populations. Regardless of fixation, they can expand into tandem repeats. Given the prevalence of MTD, we propose that they may drive the evolution of the majority of tandem repeats.

A clear example is provided by a recent experimental evolution study in *Pseudomonas fluorescens*, demonstrating complete tandem repeat evolution under fluctuating conditions (19). In this study, a 7-bp duplication with 4-bp microhomology disrupted the reading frame at the *pflu0185* locus, switching cells between non-cellulosic (CEL^−^) and cellulose-producing (CEL^+^) phenotypes. Subsequent expansions allowed rapid phenotype switching through frame-restoring contractions or expansions (Figure S6). Similarly, the highly variable MTDs identified in *S. cerevisiae* (Figure 4C) likely represent active TR evolution driven by unknown selective pressures.

By creating localized hypermutability, MTDs generate “evolvability hotspots” where functional modularity (e.g., c-di-GMP sensing in *P. fluorescens* or rapamycin resistance in *S. pombe* (6, 19)) and tandem repeat architecture synergistically enhance survival in dynamic environments. This conserved mechanism positions MTDs as genomic accelerants that balance mutational potential with phenotypic adaptability, offering a plausible strategy for tandem repeat evolution.

### Selective retention of MTDs as adaptive genetic modules

The fixation of MTDs within populations or species suggests selective advantages in fluctuating environments, such as periodic exposure to stressors like drugs or nutrient deprivation. Many fixed MTDs occur within or near coding regions, influencing gene dosage and protein functionality and significantly affecting phenotype. The intrinsic instability of these genetic elements may itself confer adaptability advantages (20).

In humans, approximately 139,000 unique tandem repeats are associated with nearby molecular phenotypes, including many known disease-risk tandem repeats, such as those linked to amyotrophic lateral sclerosis. Furthermore, around 18,700 tandem repeats have been identified as potential causal variants (21).

On the other hand, the recurrence of *de novo* MTDs (that are not fixed) in microbial populations may provide selective advantages under specific environmental pressures. While large effective population sizes in microorganisms reduce the fixation of neutral mutations by genetic drift (22), the recurrent detection of MTDs in stress-responsive loci, such as rapamycin/caffeine resistance in *S. pombe* (6) and nitrogen starvation adaptation in *S. cerevisiae* (23), suggests positive selection drives their maintenance.

Among the eighteen overlapping loci between *de novo* TDs and variable TDs in *S. cerevisiae*, five displayed multiple-copy alleles, with 66.7% (12/18) located within or proximal to coding regions. Notably, one specific locus was directly positioned at the origin of replication (Table S5). This functional enrichment implies that MTDs may fine-tune gene dosage to optimize stress responses, for example, by transiently amplifying transporters or regulators under nutrient limitation, while their inherent reversibility minimizes fitness costs upon environmental shifts.

### Implications for human health and diseases

MTDs also appear to be linked to human diseases, notably cancers. Most tandem duplications in the human genome exhibit microhomology arms, mirroring microbial patterns. This evolutionary conservation suggests functional significance. Given the reversibility of MTDs, their abundance in cancer-associated mutations highlights the need for further investigation into their role in tumor biology. Could the reversibility of disease-associated MTDs be a key factor in disease progression? How does a reversible pathogenic mutation contribute to the heterogeneity of tumors? Additionally, exploring whether targeting this reversibility offers therapeutic benefits, especially in cancers with complex genomic profiles, could provide promising avenues for future treatments.

### Mechanistic insights into MTD formation

In *cis*, microhomology arms play key roles in tandem duplication formation. Coding regions may impose selective constraints on mutations. Consistently, the proportion of TDs without MHAs (0 bp) was lower within coding regions than the genome-wide average in fungi and bacteria (Figure S7, S8), suggesting MHAs preferentially facilitate TD formation in these regions. This preference may be due to MHAs enhancing precision during duplication events.

To characterize MHA sequence features, we analyzed single-population deep sequencing data from 15 microbial species spanning bacterial, fungal, archaeal, and viral taxa. While our intra-population analysis revealed lineage-specific variation in MHA GC content relative to whole-genome baselines (Figure S9), a conserved pattern emerged: *de novo* TD sites consistently exhibited a positive correlation between MHA length and GC content across all examined taxa (Figure S10). This finding aligns with prior studies of MHA features in *S. pombe* and *E. coli* (6), where longer GC-rich MHAs demonstrated substantially higher MTD propensity. Specifically, long GC-rich MHAs are 1,000-fold more likely to generate tandem duplications than short AT-rich MHAs.

Our data indicate preferential retention of MTDs at genomic locations, suggesting they serve as dynamic generators of genetic variation useful in adaptive evolution. The persistence of duplications across generations, facilitated by MHAs, contributes to their evolutionary dominance.

In *trans*, errors in DNA replication and repair can create environments conducive to the emergence of tandem duplications, as suggested by the increased occurrence of MTDs in *rad27*Δ mutants. Errors in Okazaki fragment processing create conditions favorable for TD generation. Future research should focus on dissecting the molecular pathways driving MTD generation, especially in various mutator strains and environmental contexts.

Since the Okazaki fragments maturation process is evolutionarily highly conserved, it is conceivable that similar mechanisms may underlie the formation of MTD mutations across diverse species (Figure 1B, Table S2). However, other molecular processes beyond replication, such as transcription, may also drive MTD formation. For example, tandem repeat expansions are promoted by elevated transcription, as demonstrated experimentally in *P. fluorescens* (19). Likewise, TR expansion commonly underlies the pathological development of multiple human neuron degenerative diseases such as Huntington’s disease, amyotrophic lateral sclerosis (ALS), and fragile X-associated tremor/ataxia syndrome (FXTAS)(24, 25). These diseases usually have a late onset at the adult stage, during which neuron cells cease to proliferate (26, 27).

## Conclusion

Our findings affirm that, across all domains of life, MTD is the dominant form of TD and significantly contributes to genetic diversity, genome evolution, and disease pathogenesis. The prevalence and reversibility of MTDs underscore their versatility as a mutational mechanism, revealing genetic plasticity even within isogenic microbial populations. Similarly, somatic cells in multicellular organisms, including humans, may exhibit similar genetic variation. A deeper understanding of MTDs—especially the mechanisms behind their formation and biological implications—could enhance our knowledge of genome evolution, may inform novel strategies for addressing genetic disorders and cancers, and further the understanding of genomic plasticity and life’s resilience.

## Acknowledgments

We thank Zhenxin Yan, Qing Li, Cunqi Ye, Jie Zhou, Weiguo Fang, and Guanghua Huang for providing microbial strains. We are also grateful to Yafei Mao, Qing Li, and Jian Zhang for their critical comments on the manuscript.

## Author contributions (CRedIT)

**Author contributions (CRediT):**
| AUTHORS | ROLE |
| --- | --- |
| C.J., X.H. | Conceptualization |
| X.F.W., X.H., C.J. | Methodology |
| X.F.W., W.G., C.J. | Software |
| X.F.W., J.Z. | Validation |
| X.F.W., W.G., C.J. | Formal analysis |
| X.F.W., X.Y.W. | Investigation |
| X.H., C.J., X.Y.W. | Resources |
| X.F.W., C.J. | Data Curation |
| X.F.W., C.J., X.H. | Writing |
| X.F.W., Y.Z., C.P., C.J., | Visualization |
| C.J., X.H. | Supervision |
| C.J., X.H. | Project administration |
| C.J., X.H. | Funding acquisition |

### Data and materials availability

All processed data and code are available at https://github.com/zjumtdpro/MTDpro. The data for this study have been deposited in the European Nucleotide Archive (ENA) at EMBL-EBI under accession number PRJEB87196 (https://www.ebi.ac.uk/ena/browser/view/PRJEB87196).

## Materials and Methods

### Strains and Culturing Conditions

All bacterial and fungal strains used in this study are listed in Supplementary Table S6. *Schizosaccharomyces pombe* (fission yeast) was cultured in YE5S liquid or solid medium (supplemented with histidine, uracil, lysine, leucine, and adenine) at 29°C. *Saccharomyces cerevisiae* (budding yeast) was grown in YPD liquid or solid medium at 29°C. The *rad27*Δ deletion strains of *S. cerevisiae* (W303-1A background: MATa ade2-1 ura3-1 his3-11,15 trp1-1 leu2-3,112 can1-100) were obtained from Shi et al. (2024) (18). Species-specific culture conditions (media compositions and growth temperatures) for all other microbial species are detailed in Supplementary Table S6. All cultures were harvested at the mid-logarithmic growth phase (optical density OD₆₀₀ of 0.5–0.8) for subsequent experiments.

### Library Preparation and Sequencing

#### Genomic DNA extraction and QC

DNA was extracted using phenol-chloroform purification. Concentration was measured by Qubit® 3.0 Fluorometer (Thermo Fisher Scientific), and integrity assessed via 1% agarose gel electrophoresis. Samples with 260/280 = 1.8-2.0 and 260/230 >2.0 were used for library construction.

#### Library construction

Libraries were prepared from 200 ng DNA using Watchmaker DNA Library Prep Kit with Fragmentation (Watchmaker Genomics; CAS:7K0019-096). DNA was enzymatically fragmented (∼350 bp), followed by end repair, A-tailing, and adapter ligation per manufacturer’s protocol. Adapters and indices were incorporated via limited-cycle PCR (8–10 cycles).

#### Sequencing

Library size distribution (300–500 bp) and absence of adapter dimers were confirmed by Fragment Analyzer (Agilent Technologies). Quantification used KAPA Library Quantification Kit (Roche) on ABI QuantStudio 12K Flex. Pooled libraries were sequenced on DNBSEQ-T7 (MGI Tech) with DNBSEQ-T7RS High Throughput Kit (FCL PE150) V3.0, generating 150-bp paired-end reads at 50× coverage. Technical replicates of *S. cerevisiae* and *E. coli* were sequenced on Illumina NovaSeq 6000 with NovaSeq S4 Reagent Kit V1.5 under identical parameters.

### Bioinformatic Analysis

#### Data preprocessing

Raw sequencing reads were quality-assessed using FastQC (v0.11.9) and processed using fastp (v0.22.0) (28) with the following parameters: quality filtering (minimum Phred score: 20), adapter trimming (auto-detection), and read filtering (minimum length: 50 bp). Quality metrics were aggregated using MultiQC (v1.9) to ensure consistent data quality across all samples.

#### Alignment

Processed reads were aligned to species-specific reference genomes (detailed in Supplementary Table S1) using bwa mem (v1.11) (29) with soft-clipping enabled (-Y flag) and default alignment parameters. Post-alignment filtering excluded supplementary alignments (samtools view -F 2048) and unmapped reads (samtools view -F 4). Alignment files were sorted using samtools sort (v0.7.17) (30) and indexed using samtools index. Alignment statistics, including genome coverage, mapping rate, and duplication rate, were calculated using samtools flagstat and bedtools genomecov (v2.30.0).

#### MTD identification

MTDs were systematically identified using a comprehensive computational pipeline. The core detection was performed using MTD-finder (available at https://github.com/carey-lab/MicroHomologyMediatedTandemDuplications).

To enhance detection sensitivity and distinguish between MTDs and NMTDs, we developed an improved pipeline (MTDfinder-pro) with the following steps:

1. INDEL calling: Genomic coverage profiles were generated using samtools mpileup (v1.15) with default parameters. Putative insertions and deletions (INDELs) were identified using VarScan (v2.3.9) (31) with the pileup2indel module (parameters: --min-var-freq 0).
2. Candidate TD screening: For each identified insertion, flanking sequences (including two repeat units downstream of the insertion site) were extracted using bedtools getfasta (v2.30.0) (32). Tandem duplication structures (1×→2× repeat expansions) were confirmed using Tandem Repeat Finder (v4.09.1) (33) with parameters optimized for short repeats (match = 2, mismatch = 7, delta = 7, PM = 80, PI = 10, minscore = 30) to detect tandem repeats ≥ 8 bp with 2× periodicity.
3. Microhomology analysis: Flanking sequences of confirmed tandem duplications were analyzed using a dynamic programming algorithm to detect MHAs. The algorithm identified the longest perfect sequence match at the breakpoint junctions. Events with MHA ≥ 1 bp were classified as MTDs, while those without detectable microhomology (MHA = 0 bp) were classified as NMTDs. The source code for microhomology analysis is available at https://github.com/zjumtdpro/MTDpro.

### Experimental Validation

#### Sanger sequencing

Genomic DNA was extracted from *S. pombe* using phenol-chloroform purification and from *E. coli* via boiling lysis (95°C for 10 min in 1× TE buffer). Target loci were amplified using PrimeSTAR® Max DNA Polymerase (Takara Bio; RR006Q) with primers listed in Table S7, in 25 μL reactions containing 12.5 μL 2× Premix, 0.3 μM primers, and 50–100 ng DNA. Thermal cycling conditions were: 98°C for 3 min; 30 cycles of 98°C for 10 s, 55°C for 15 s, and 72°C for 30 s/kb; final extension at 72°C for 5 min. Purified PCR products (ExoSAP-IT™, Thermo Fisher) underwent bidirectional Sanger sequencing, and chromatograms were analyzed using SnapGene Viewer (v5.1).

#### Single-Cell Whole-Genome Sequencing (*S. cerevisiae*)

Single-cell sequencing was performed using the MobiNova®-M1 microfluidic platform (MobiDrop) following manufacturer protocols with modifications(34, 35). Mid-log phase *S. cerevisiae* cells were washed with PBS, resuspended to 4×10^8^ cells/mL, and mixed with 15% (v/v) OptiPrep density gradient medium (Sigma-Aldrich D1556) to prevent aggregation. For lysis, 100 μL cell suspension and 240 μL lysis buffer (400 mM KOH, 100 mM DTT, 10 mM EDTA, 2% Tween-20) were partitioned into 75-μm droplets. Thermal lysis proceeded sequentially: 37°C (30 min), 75°C (10 min), and 95°C (5 min), followed by rapid cooling to 4°C. Whole-genome amplification used φ29 polymerase-mediated multiple displacement amplification (MDA) in 120 μL reactions (30°C, 8 h; 65°C, 10 min inactivation). Libraries were prepared by: (1) Tagmentation with 108 μL Nextera mix (55°C, 10 min); (2) Barcoding via dual-indexed primers in 308 μL PCR mix (72°C, 4 min; 98°C, 30 s; 10 cycles of 98°C/7 s, 60°C/30 s, 72°C/40 s; 72°C, 5 min). Post-barcoding, droplets were disrupted with 1H,1H,2H-perfluoro-1-octanol, and DNA purified using SPRIselect beads (0.7× ratio). Bioinformatics analysis followed established pipelines(34), with quality control via BUSCO (36, 37).

### Multi-level Evolutionary Analysis

#### Intra-species Level Analysis

Comprehensive microbial genomic datasets were retrieved from NCBI GenBank and RefSeq databases (Release 2021.07, accessed July 6, 2021). Species selection was based on the availability of a high-quality reference genome and at least two additional high-quality isolate genomes. Whole-genome alignment was performed using MUGSY (v1.2.3) with default parameters (38) to align isolate genomes to their respective reference genomes.

Insertions ≥ 8 bp were identified from the resulting Multiple Alignment Format (MAF) files using a custom Python (v3.9.21) script (available at https://github.com/zjumtdpro/MTDpro). TDs and their associated MHAs were annotated using the MTDfinder-pro pipeline described above. For statistical robustness, bacteria and fungi with ≥ 20 validated TD loci and archaea with ≥ 10 validated TD loci were retained for further analysis.

For the identification of 3x copy number variants, characteristic 3x signature sequences were generated for each identified TD locus by duplicating the corresponding insertion sequence. These signature sequences were systematically cross-referenced against the complete set of insertion mutations for each species through coordinate-based matching and sequence alignment. Matches indicated the independent occurrence of 3x alleles at TD loci, and the frequency of such occurrences was recorded.

#### Species Level Analysis

A representative set of 1,000 genomes per kingdom (fungi, protozoa, bacteria, archaea) was randomly selected from the same NCBI database. Reference genomes were analyzed using TRF (v4.09.1) with parameters optimized to detect tandem repeats ≥ 8 bp with a minimum score of 30. For loci with 100% identity between repeat units, MHA length was calculated using custom awk scripts. For loci with <100% identity between repeat units, the proportion of positions with exactly two copies (indicative of tandem duplication without microhomology) was determined.

#### Human Dataset Analysis

1. Reference genome characterization: TRs and TDs in GRCh38/hg38 were identified using TRF (v4.09.1, min-score=30). For perfect repeats (≥8 bp, 100% unit identity), MHA lengths were calculated with custom awk scripts.
2. Individual analysis: Variable tandem repeats from the Eslami et al. (2021) (39) dataset were categorized as tandem duplications (2x) or higher-order tandem repeats (≥3x), with MHA proportions quantified for each category using custom bioinformatic scripts.
3. Somatic-germline mutation profiling: Whole-genome sequencing data from Moore et al. (2021) (10) encompassing 389 laser-microdissected histological patches (200-1,000 cells each) across 29 tissues from three individuals (27× median coverage) were re-analyzed, with *de novo* indel calling followed by tandem duplication identification via TRF (100% unit match required) and subsequent microhomology arm length determination.
4. Disease database analysis: Disease-associated insertion/deletion mutations (≥4 bp) were curated from COSMIC v99 (GRCh37 variants: Cosmic_MutantCensus_v99_GRCh37, Cosmic_NonCodingVariants_v99_GRCh37) and ClinVar (January 5, 2024 release), excluding low-complexity sequences. Mutations were classified through sequential analysis: variants within annotated tandem repeat regions were assessed by comparing indel sequences to consensus repeat units to determine copy number alterations, while variants in non-repetitive regions underwent *de novo* tandem repeat identification using TRF (v4.09.1), retaining only events with 100% identity between adjacent copies for TD confirmation and MHA annotation.

### Statistical Analysis and Visualization

Permutation tests (one-tailed, 100,000 simulations) were performed to assess deviations from random expectations for MHAs. Group-level *p-values* were calculated using a binomial model (expected probabilities: 75% for MHA=0 bp, 25% for MHA ≥ 1 bp, 6.25% for MHA ≥ 2 bp, 1.56% for MHA ≥ 3 bp). Analyses were implemented in Python (v3.9.21), and visualizations were generated using ggplot2 (v3.5.1) and circlize (v0.4.16) (40) in R (v4.3.2). The phylogenetic tree is generated and visualized using pctax (v0.1.3).

---

**Figure S1.**
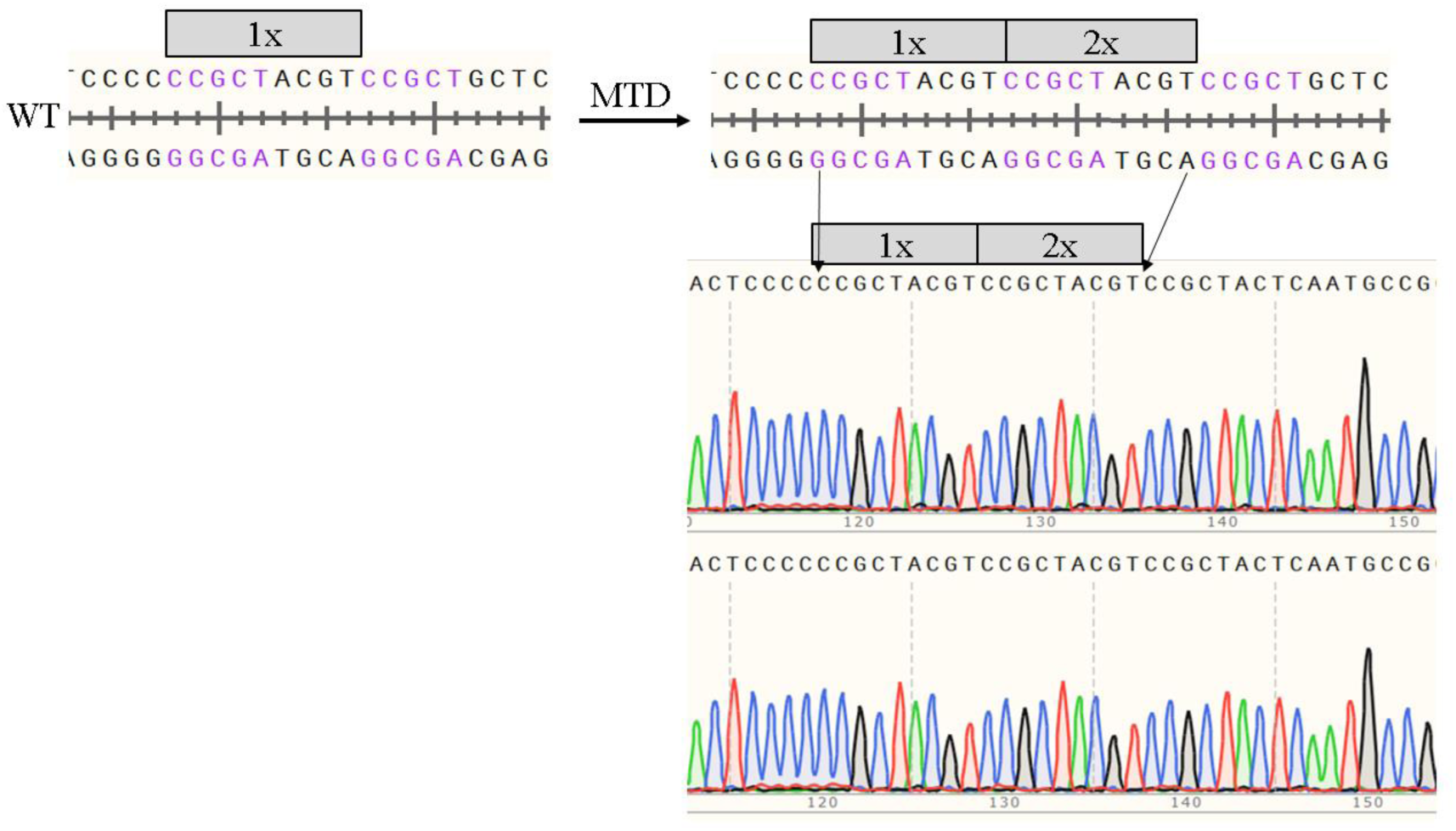
Validation of MTD loci by sanger sequencing. The MTD site is located at NC_000913.3: 2029728–2029736 in *E. coli*. The left panel displays the reference sequence, while the right panel shows the sequence following MTD formation, with microhomology arms highlighted in purple. Below, Sanger sequencing results for two single clones are presented. Due to space constraints, additional MTD loci are not shown here; detailed information on species, loci, and primers can be found in Table S7.

**Figure S2.**
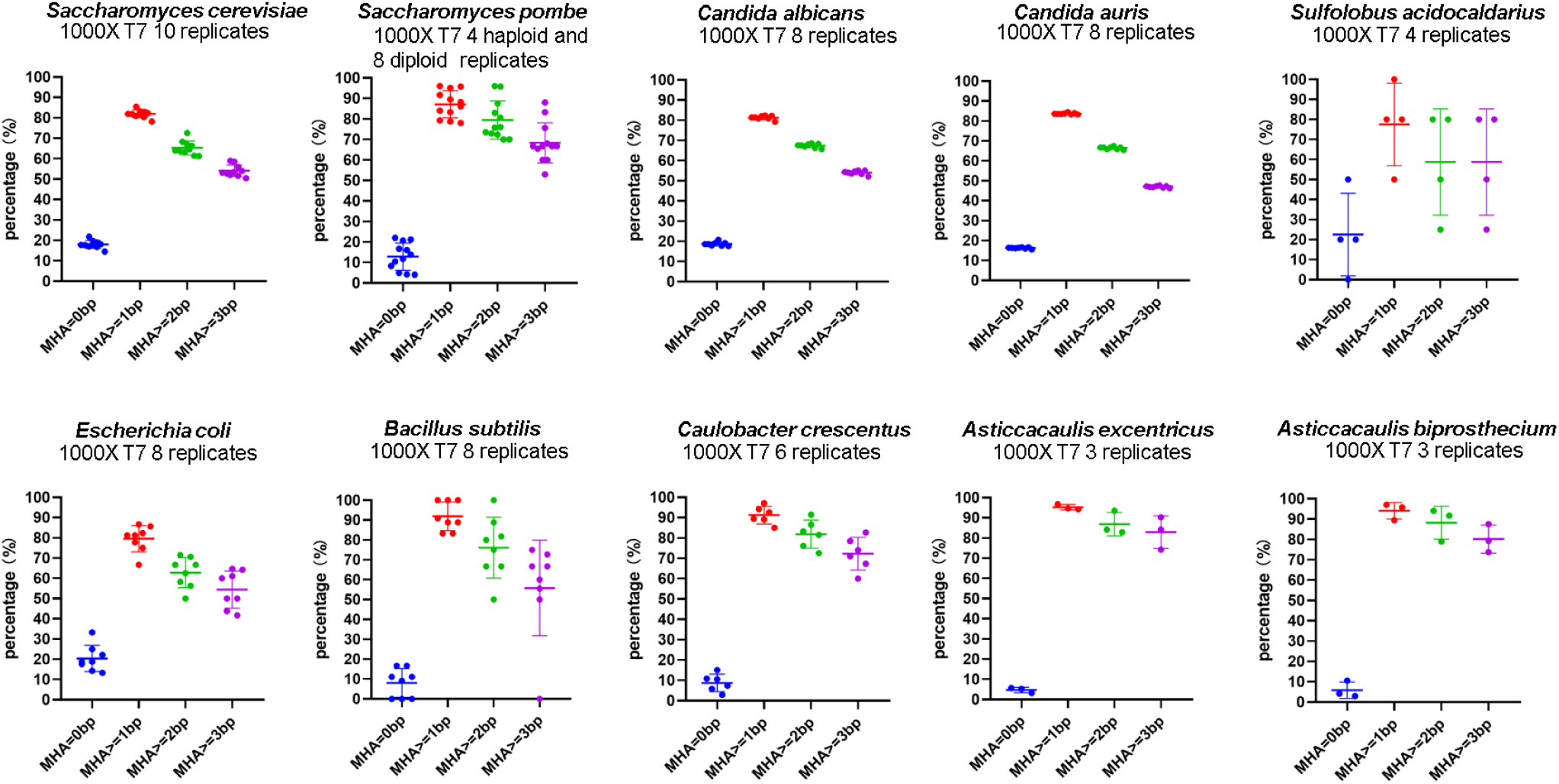
Proportions of TDs with MHA in single-population deep sequencing data from ten species. Each point represents a sequencing replicate. The data were generated using the DNBSEQ-T7 sequencing platform, with each sample sequenced at a depth of 1,000×.

**Figure S3.**
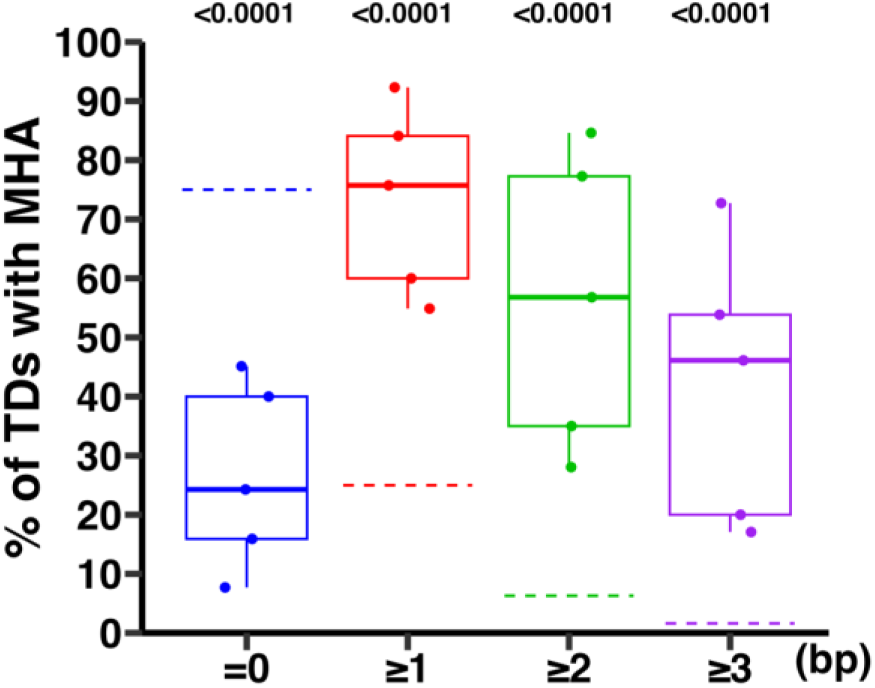
Proportions of variable TDs with MHA at the intra-species level of archaea. Each point represents a species, and the dashed line represents the random expected value. *P-values* were calculated using permutation tests (one-tailed, 100,000 simulations) against theoretical expectations. Data are presented for species with at least 10 TD loci.

**Figure S4.**
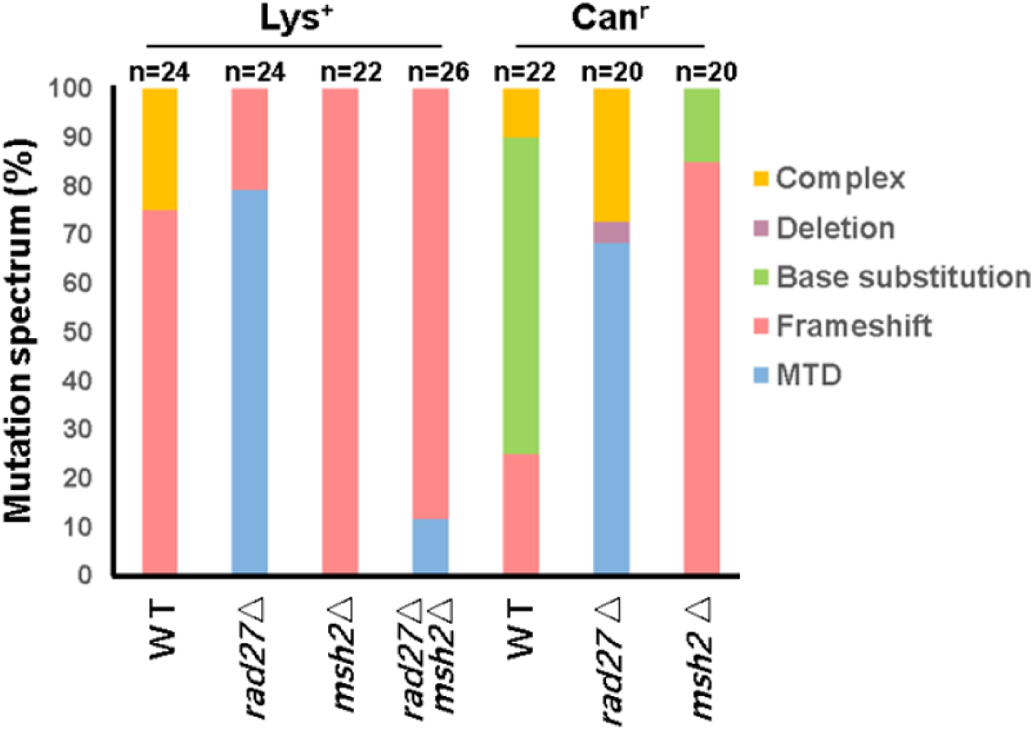
Mutation spectrum of Lys^+^ and Can^r^ revertants in *rad27*Δ and *msh2*Δ mutant strains of *S. cerevisiae*. The identified loci are Lys^+^ (left) and Can^r^ (right). Mutation types are color-coded, with “n” representing the sample size.

**Figure S5.**
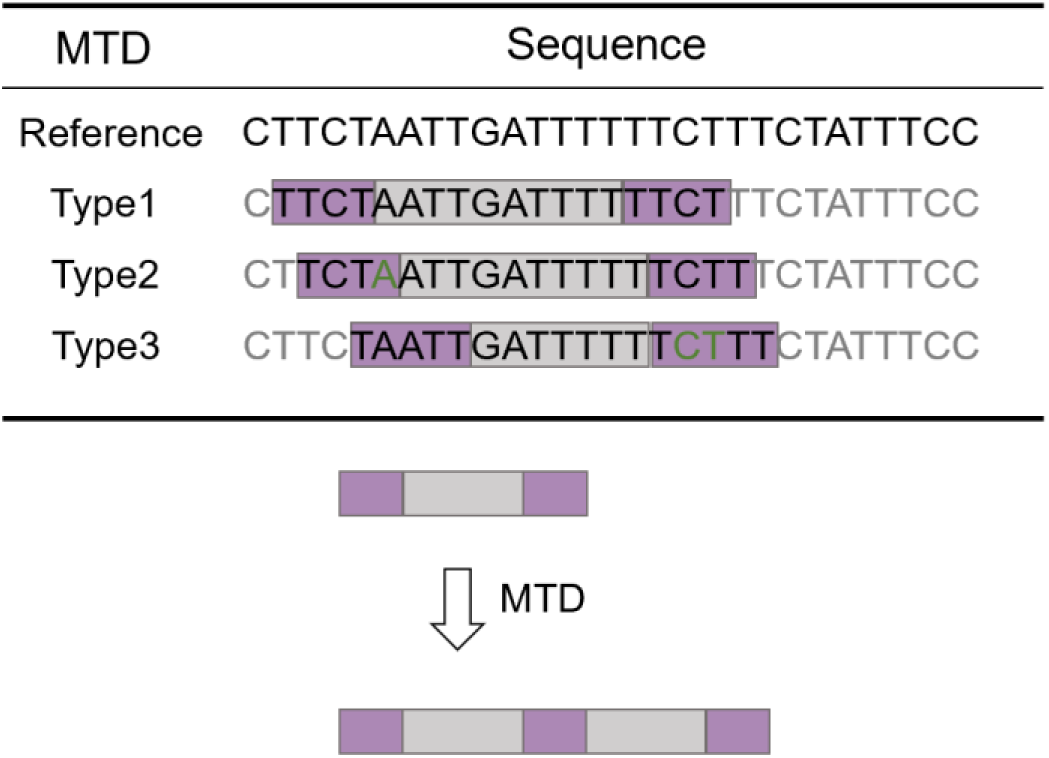
Representative MTD clusters with overlapping coordinates. (Top) Reference sequence with annotated MTD types (Type 1–3). Purple: MHAs; green: mismatched bases. (Bottom) Schematic model of MTD formation.

**Figure S6.**
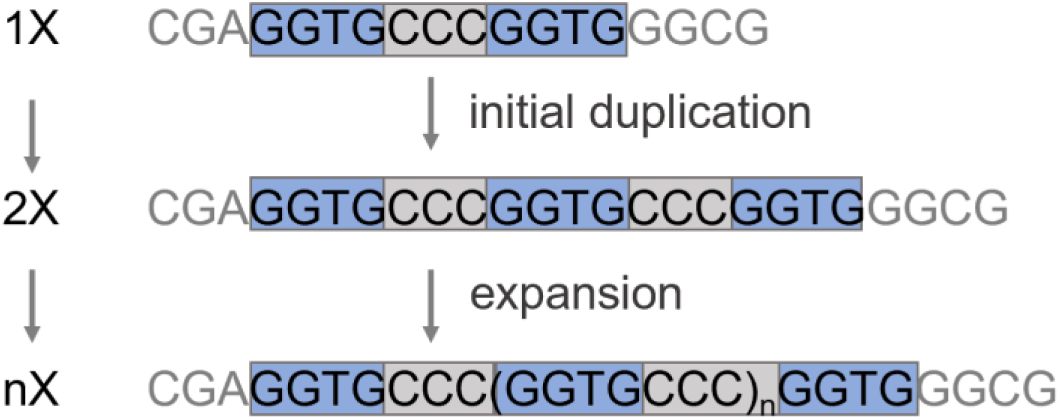
Schematic of MTD-driven evolvability in *Pseudomonas fluorescens*. The DNA sequence depicts the MTD locus within *pflu0185*. Blue bases (GGTG) highlight the microhomology arms.

**Figure S7.**
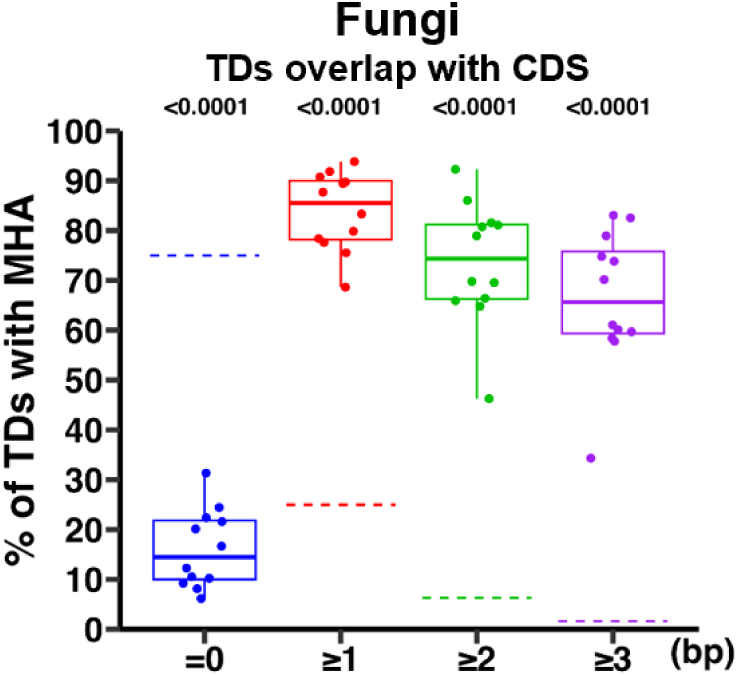
Proportions of TDs located within CDS (Coding Sequence) with MHA of fungi. Each point represents a species, and the dashed line represents the random expected value. *P-values* were calculated using permutation tests (one-tailed, 100,000 simulations) against theoretical expectations. Data are presented for species with at least 20 TD loci.

**Figure S8.**
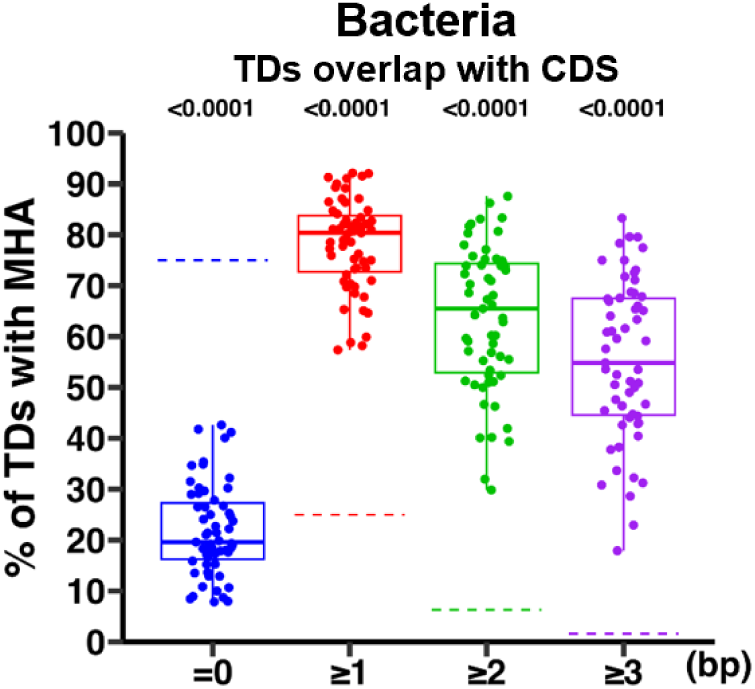
Proportions of TDs located within CDS with MHA of bacteria. Each point represents a species, and the dashed line represents the random expected value. *P-values* were calculated using permutation tests (one-tailed, 100,000 simulations) against theoretical expectations. Data are presented for species with at least 20 TD loci.

**Figure S9.**
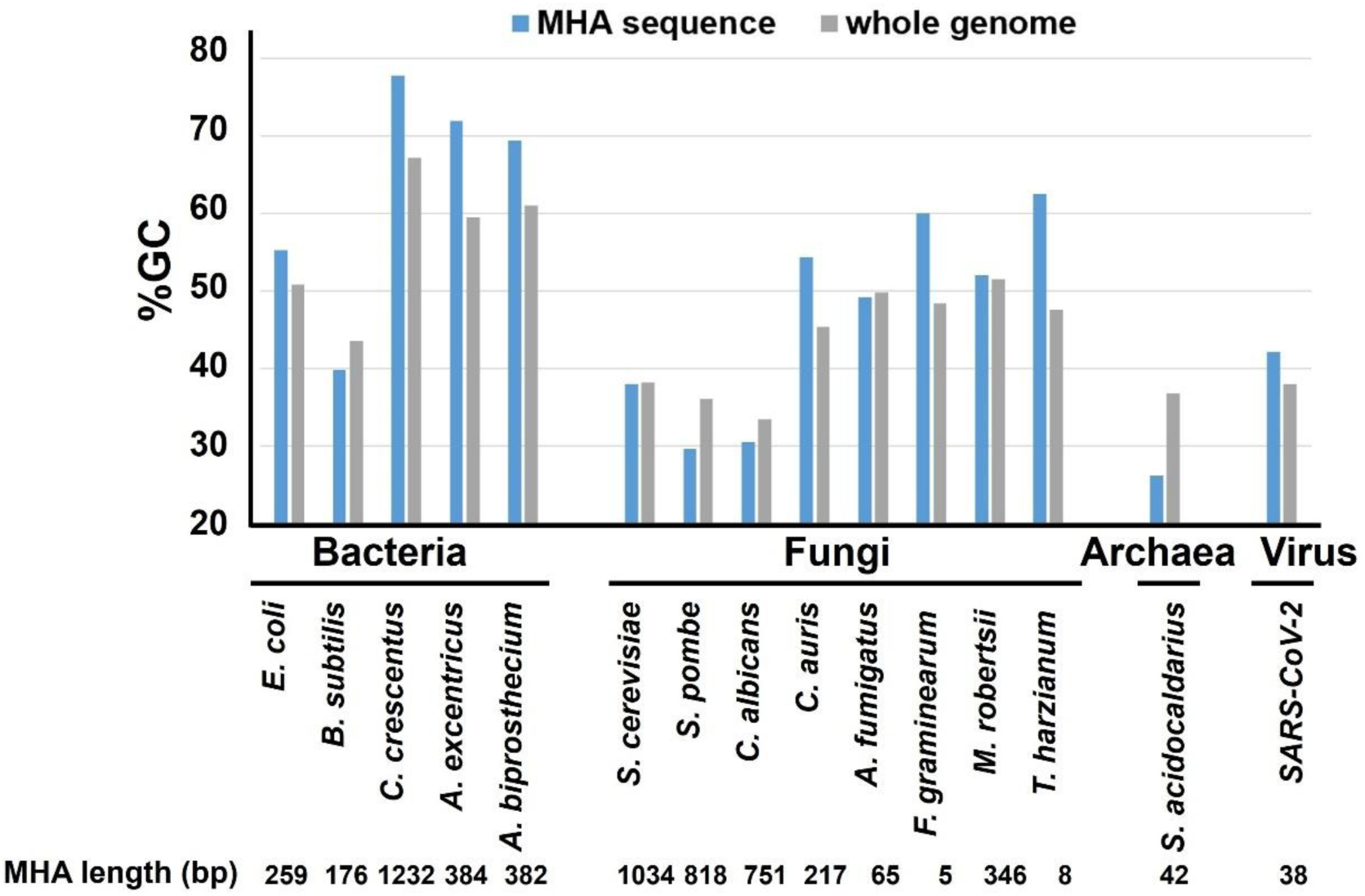
Comparative analysis of GC content in MHAs versus whole genomes across taxonomic groups. Blue bars represent %GC of MHA sequences; gray bars indicate whole-genome average GC%. Analyses include bacterial, fungal, archaeal, and viral species. Lower panel: total MHA length for corresponding species.

**Figure S10.**
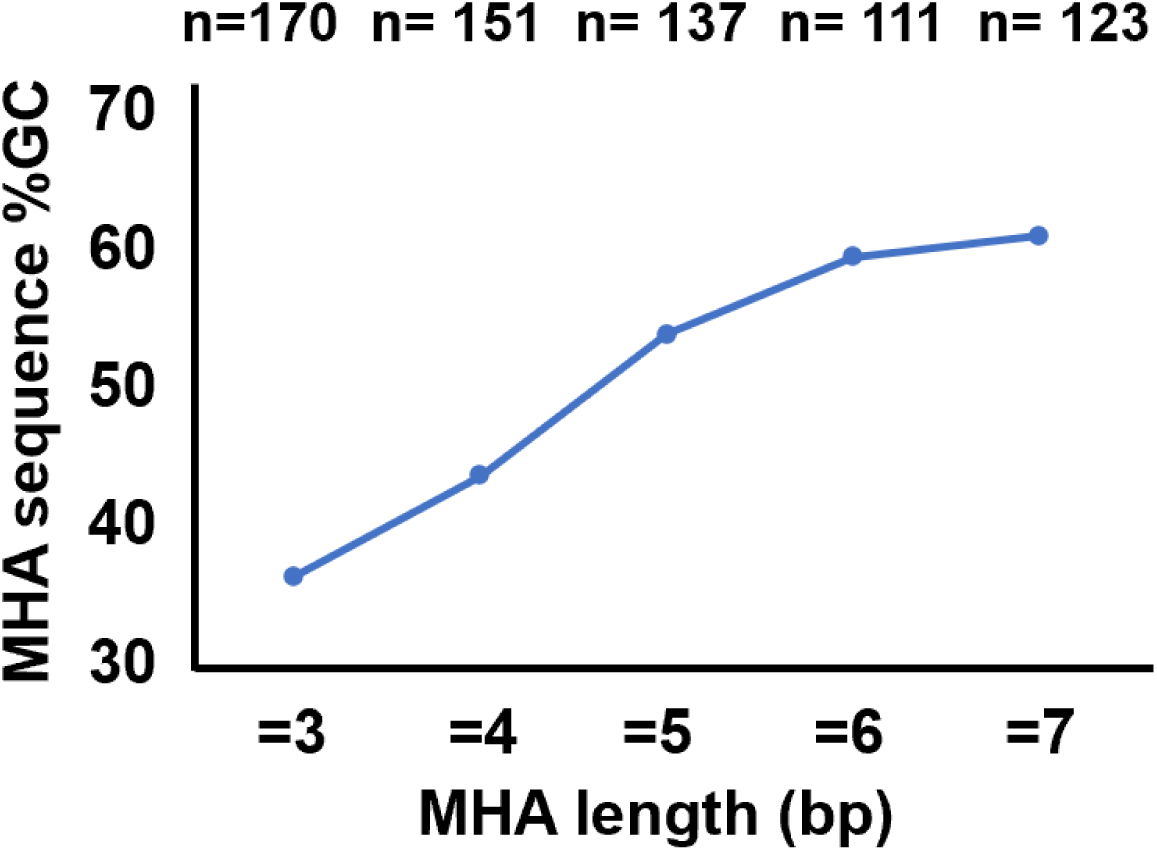
Correlation between MHA sequence length and GC content at the intra-population level. Data derived from 15microbial species. Top panel: number of MTD loci (n) for each MHA length category.

Supplementary_tables are in an Excel file.

